# Reduced Monocyte Proportions and Responsiveness in Convalescent COVID-19 Patients

**DOI:** 10.1101/2023.10.25.563806

**Authors:** Eugene V. Ravkov, Elizabeth S.C.P. Williams, Mark Elgort, Adam P. Barker, Vicente Planelles, Adam M. Spivak, Julio C. Delgado, Leo Lin, Timothy M. Hanley

## Abstract

The clinical manifestations of acute severe acute respiratory syndrome coronavirus type 2 (SARS-CoV-2) infection and COVID-19 suggest a dysregulation of the host immune response that leads to inflammation, thrombosis, and organ dysfunction. It is less clear whether these dysregulated processes persist during the convalescent phase of disease or during long COVID. We investigated the effects of SARS-CoV-2 infection on the proportions of classical, intermediate, and non-classical monocytes, their activation status, and their functional properties in convalescent COVID-19 patients and uninfected control subjects. We found that the percentage of total monocytes was decreased in convalescent COVID-19 patients compared to uninfected controls. This was due to decreased intermediate and non-classical monocytes. Classical monocytes from convalescent COVID-19 patients demonstrated a decrease in activation markers, such as CD56, in response to stimulation with bacterial lipopolysaccharide (LPS). In addition, classical monocytes from convalescent COVID-19 patients showed decreased expression of CD142 (tissue factor), which can initiate the extrinsic coagulation cascade, in response to LPS stimulation. Finally, we found that monocytes from convalescent COVID-19 patients produced less TNF-α and IL-6 in response to LPS stimulation, than those from uninfected controls. In conclusion, SARS-CoV-2 infection exhibits a clear effect on the relative proportions of monocyte subsets, the activation status of classical monocytes, and proinflammatory cytokine production that persists during the convalescent phase of disease.

## INTRODUCTION

Severe acute respiratory syndrome coronavirus type 2 (SARS-CoV-2) is the causative agent of coronavirus infectious disease 19 (COVID-19) As of August 2023, it has affected over 770 million people, resulting in nearly 7 million deaths worldwide (WHO Coronavirus (COVID) dashboard). The symptoms of the disease vary from mild to severe, depending on age, gender, physical health, and other host factors. Notably, a dysregulated host immune response is observed in COVID-19 patients, leading to complications like hyperinflammation, thrombosis, and organ damage ^1, 2^. Cytokine storm, characterized by elevated levels of circulating cytokines and hyperactivation of immune cells ^3, 4^, and coagulopathy, an alteration in the normal blood clotting process ^5, 6^, are the two major mechanisms contributing to COVID-19 pathogenesis. Myeloid cells, including circulating monocytes and tissue macrophages, are pivotal in mediating these pathologies.

Alterations in myeloid cells, particularly monocytes, are evident during acute COVID-19 infection. Specifically, total monocyte counts are decreased, especially in severe disease ^7, 8^. In addition, multiple studies demonstrate differences in the proportions of classical (CD14^+^ CD16^-^), intermediate (CD14^+^ CD16^+^), and non-classical monocytes (CD14^lo^ CD16^+^), with a notable decrease in circulating non-classical monocytes in moderate and severe COVID-19 ^9–12^.

Phenotypic changes in monocytes in patients with moderate and severe COVID-19 infection have also been described, including increased CD83 expression ^9^ and decreased HLA-DR expression ^9–13^. During acute infection, monocytes and tissue macrophages are characterized by increased activation marker and inflammatory gene expression as measured by single cell transcriptomics ^14, 15^. This increase in activated monocytes and macrophages corresponds to increased levels of chemokines and cytokines ^15^, including TNF-α and IL-1β, and increased interferon-stimulated gene (ISG) expression ^16–18^. Together, these studies suggest that SARS-CoV-2 infection induces myeloid cells to generate a hyperinflammatory state similar to cytokine storm during acute infection.

The fate of monocytes during the convalescent phase of COVID-19 is less well understood. In some studies, patients who fully recovered from acute COVID-19 infection were found to have increased circulating non-classical monocytes and decreased classical monocytes during convalescence ^19–21^. In addition, these studies showed that monocytes in convalescent individuals display increased levels of inflammatory genes ^21–24^, antigen presentation molecules, including HLA-DQA and HLA-DPA ^20^, and the activation marker CD169 ^21^. However, other studies have demonstrated a decrease in monocyte percentages in convalescent individuals including those who had severe disease ^25, 26^. While a growing body of research has shed light on monocyte behavior during the convalescent phase, contrasting findings underscore the need for more detailed studies to unravel these complexities.

Similar findings have been seen with long COVID, a heterogeneous syndrome characterized by persistent significant health issues lasting, recurring, or developing three months after initial infection ^27–30^. Although the mechanisms driving long COVID are poorly understood ^31^, myeloid cells are thought to play a significant role. A number of studies have demonstrated that patients experiencing long COVID have persistently elevated levels of IFN-β, IFN-λ1, IFN-γ, TNF-α, IL-1β, IL-6, IL-10, IL-12, and IL-17 ^32–35^; however, more recent studies have demonstrated that long COVID is associated with cytokine deficiencies ^36^. In addition, Phetsouphanh et al. described elevated numbers of circulating activated monocytes and plasmacytoid dendritic cells (pDCs) ^35^. Despite these studies, the contribution of myeloid cells in long COVID, much like in the convalescent phase of the disease, are not well understood.

To further clarify the role and behavior of monocytes post-COVID, our study used a comprehensive approach involving multiparameter flow cytometry and functional assays. We focused on the phenotypic and functional characteristics of monocytes in the circulating blood from convalescent COVID-19 patients. Specifically, we examined the proportions of classical, intermediate, and non-classical monocytes, evaluated their activation status in response to bacterial lipopolysaccharide (LPS), and examined their ability to produce inflammatory cytokines. Our studies revealed several phenotypic and functional alterations in monocytes during the convalescent phase of SARS-CoV-2 infection.

## MATERIALS AND METHODS

### Study subjects

We obtained donor blood samples from individuals who were recruited under University of Utah Institutional Review Board (IRB) protocol 131664. These individuals were recruited from Salt Lake City, UT and the surrounding metropolitan area between May of 2020 and December of 2021. Informed consent was obtained from all individual participants included in the study. The de-identified specimens were blinded during collection and analysis. Whole blood samples were drawn from 18 convalescent COVID-19 patients and 30 healthy individuals in parallel. The clinical samples were obtained from convalescent patients who exhibited a mild form of the disease and did not require hospitalization. For convalescent patients, specimens were collected approximately two to four weeks after disease onset. Donors reported no symptoms at the time of sample collection. The COVID-19 patients ranged in age from 19 to 65 years old (43.3 y/o mean), representing different races and genders. The healthy control individuals ranged in age from 19 to 71 years old (38.5 y/o mean), representing different races and genders (Table 1). SARS-CoV-2 infection was assessed by one or more immunologic (Abbott Architect SARS-CoV-2 IgG, Euroimmun Anti-SARS-CoV-2 ELISA, or Siemens SARS-CoV-2 IgG) {Rychert, 2021, 33064790} or molecular assays.

**Table 1.**

| Patient Characteristics (n=49) | Control (n=31) | COVID-19 (n=18) | P value |
| --- | --- | --- | --- |
| Age, y |  |  |  |
| Mean | 38.5 | 43.3 | 0.3098<sup>a</sup> |
| Median | 36 | 36 |  |
| Range | 19-71 | 19-65 |  |
| Sex, No. (%) |  |  |  |
| Male | 12 (39%) | 12 (67%) | 0.0792<sup>b</sup> |
| Female | 19 (61%) | 6 (33%) |  |
| Race, No. (%) |  |  |  |
| White | 27 (87%) | 15 (83%) | >0.9999<sup>c</sup> |
| Black or African American | 0 (0%) | 0 (0%) |  |
| Asian | 2 (6.8%) | 2 (11%) |  |
| More than one race | 0 (0%) | 1 (6%) |  |
| Unknown/Not reported | 2 (6.8%) | 0 (0%) |  |
| Ethnicity |  |  |  |
| Non-Hispanic | 21 (68%) | 13 (72%) | >0.9999<sup>d</sup> |
| Hispanic | 5 (16%) | 4 (22%) |  |
| Unknown/Not reported | 5 (16%) | 1 (6%) |  |

<sup>a</sup>Data analyzed using unpaired t-test with Welch's correction
<sup>b</sup>Data analyzed using Fisher's exact test (male vs. female)
<sup>c</sup>Data analyzed using Fisher's exact test (white vs. non-white)
<sup>d</sup>Data analyzed using Fisher's exact test (Non-Hispanic vs. Hispanic)

### Blood samples

Fifteen mL of whole blood was collected by phlebotomy-certified research staff into two BD Vacutainer EDTA Additive Blood Collection Tubes. Tubes were gently inverted eight times to mix the blood and EDTA and were then centrifuged at 150 x *g* for 20 minutes at room temperature. Peripheral blood mononuclear cells (PBMCs) were isolated by Ficoll density gradient (Histopaque-1077, Sigma), and were cryopreserved in 1 ml aliquots in 90% complete culture media (RPMI 1640 with 10% Fetal Bovine Serum (FBS)) and 10% dimethyl sulfoxide (DMSO) in sterile cryovials. PBMCs were stored in liquid nitrogen prior to use.

### Isolation of peripheral blood monocytes

Total monocytes were isolated by negative selection using the EasySep Human Monocyte Enrichment Kit without CD16 Depletion (StemCell Technologies) using the manufacturer’s instructions. Briefly, PBMCs were resuspended at 5 × 10^7^ cells/mL in isolation buffer (PBS with 2% FBS and 1 mM EDTA) and incubated with the enrichment cocktail (50 μL/mL) at 4°C for 10 minutes. Next, vortexed magnetic particles were added, and the sample was mixed and incubated at 4°C for 5 minutes. Isolation media was then added to a final volume of 2.5 mL and the sample was placed in an EasySep magnet at room temperature for 2.5 minutes. The enriched cell suspension (non-monocytes) was poured off into a new tube, and the remaining cells (monocytes) were washed x3 with isolation media. The tube was then removed from the magnet and purified monocytes were decanted into a new tube for use.

### Flow Cytometry

Two antibody panels were used in this study. The first antibody panel was designed to identify total monocytes, classical monocytes, non-classical monocytes, intermediate monocytes, B cells, T cells, NK cells, conventional or myeloid dendritic cells (mDCs), and plasmacytoid dendritic cells (pDCs). The cell subsets were identified using the following antibodies: CD64 PE-Cy7 (Cat. # B06025, Beckman Coulter); CD45 PerCP (Cat. # 340665, BD Biosciences); Fixable Viability eFluorTM 506 (Cat. # 65-0866-18, Thermo Fisher Scientific); CD14 V450 (Cat. # 655114, BD Biosciences); CD16 FITC (Cat. # 656147, BD Biosciences); CD3 APC R700 (Cat. # 659110, BD Biosciences); CD19 BV786 (Cat. # 563325, BD Biosciences); HLA-DR APC-H7 (Cat. # 641402, BD Biosciences); CD123 PE (Cat. # 649453, BD Biosciences); CD11b APC (Cat. # 340936, BD Biosciences); CD11c BV605 (Cat. # 663799, BD Biosciences). The gating strategy started with a time gate, followed by exclusion of dead cells (CD45 vs Fixable Viability Dye), and exclusion of doublets (FSC-A vs FSC-H). Monocytes were then selected as CD64+CD3-CD19-gated cells. The initially identified monocytes were then refined as HLA-DR+, CD123-, CD11b^hi^, and CD11c^hi^ cells to exclude DCs. Monocytes were then analyzed for classical, non-classical, and intermediate monocytes by comparing CD14 and CD16 expression. The mDCs were identified from the total WBC population as CD3-, CD19-, HLA-DR+, CD123-, CD11c+, CD11b-, CD14-, and CD16-cells. The pDCs were identified as CD14-, CD16-, HLA-DR-, and CD123+ cells. In addition, our gating also included identification of B cells, T cells, and NK cells as CD45+CD64-CD19+, CD45+CD64-CD3+, and CD45+CD64-CD19-CD3-CD16+ cells, respectively.

The second antibody panel was designed examine the major monocyte subsets (classical, non-classical, and intermediate) and expression of their activation markers, including a marker associated with the extrinsic coagulation pathway (CD142). The antibody panel was used for staining of both short time-stimulated (2-3 hours) and unstimulated samples. The stimulation was carried out at 37°C in CO2 incubator in the presence of 1.0 μg/mL *E. coli* liposaccharide (LPS, Invivogen) in RPMI 1640 media supplemented with 10% heat-inactivated Fetal Bovine Serum (Gibco Fetal Bovine Serum, Cat. No. 26140087), L-glutamine (200 mM L-Glutamine, ThermoFisher Scientific Cat. No. 25030149), penicillin-streptomycin (Gibco Penicillin-Streptomycin, Cat. No. 15140148). After incubation with and without LPS, cells were washed with 1% FBS in PBS antibody staining buffer and stained with the following antibodies: CD64 PE-Cy7 (Cat. # B06025, Beckman Coulter); CD45 PerCP (Cat. # 340665, BD Biosciences); Fixable Viability eFluorTM 506 (Cat. # 65-0866-18, Thermo Fisher Scientific); CD14 V450 (Cat. # 655114, BD Biosciences); CD16 FITC (Cat. # 656147, BD Biosciences); CD4 APC-H7 (Cat. # 641407, BD Biosciences); CD56 APC R700 (Cat. # 657887, BD Biosciences); CD69 APC (Cat. # 654663, BD Biosciences); CD83 BV786 (Cat. # 565336, BD Biosciences); CD86 BV605 (Cat. # 562999, BD Biosciences); CD142 PE (Cat. # 550312, BD Biosciences). The gating strategy started with a time gate, followed by exclusion of dead cells (CD45 vs Fixable Viability Dye), and exclusion of doublets (FSC-A vs FSC-H). Total monocytes were subsequently selected as CD64+ cells. The identified monocytes were then analyzed for classical, non-classical, and intermediate monocytes by CD14 and CD16 expression. Each cell subset was examined for the expression of CD4, CD56, CD69, CD83, and CD86 activation markers and the marker for the extrinsic coagulation pathway (CD142).

### Determination of cytokine concentration

Isolated monocytes were plated at 1×10^5^ cells/well in 96-well plates in 200 μL of complete culture medium (RPMI 1640 with 10% FBS, 100 U/mL penicillin, 100 μg/mL streptomycin, and 0.29 mg/mL L-glutamine). Monocytes were treated with *E. coli* LPS (Invivogen) at 100 ng/mL. After 18 hours, the plates were briefly centrifuged, and cell-free supernatant was harvested for analysis. TNFα and IL-6 were measured in culture supernatant by ELISA (R&D Systems) according to the manufacturer’s instructions.

### Statistical analysis

Demographic data of donors were analyzed for statistical significance using either unpaired t-test with Welch’s correction or Fisher’s exact test. Flow cytometry data and individual cytokine values for uninfected and convalescent COVID-19 (unpaired) specimens were analyzed for statistical significance using Mann-Whitney tests. Flow cytometry data and individual cytokine values for untreated and LPS-treated (paired) specimens were analyzed for statistical significance using Wilcoxon rank tests. The level of significance was determined as p < 0.05. Statistical analyses and graphic presentation were performed using Prism 9.01 software (GraphPad Software, Inc., San Diego, California).

## RESULTS

### Study population description

Peripheral blood specimens were obtained from uninfected individuals and from convalescent patients who exhibited a mild form of the disease. Specimens were collected between May 2020 and January 2021, during the initial SARS-CoV-2 outbreak in the United States. The median age of the participants was 36 years (range 19-65) for uninfected subjects (n=31) and 42.5 years (range 19-71) for convalescent COVID-19 subjects (n=18). None of the participants had received a COVID vaccine at the time of sample collection. All individuals from the convalescent COVID-19 population had mild symptoms, including fever, headache, rhinorrhea, sore throat, cough, fatigue, and/or myalgias, which did not require medical intervention or hospitalization during illness. Their symptoms resolved within two weeks of disease onset and specimens were collected approximately two to four weeks after disease onset. The convalescent patients and uninfected donors reported no symptoms at the time of sample collection. Participant characteristics are listed in Table 1.

### The proportion of circulating monocytes is decreased in convalescent COVID-19 patients

To determine the effect of COVID-19 infection on blood monocytes during the convalescent stage of disease, peripheral blood mononuclear cells were isolated from convalescent COVID-19 patients and uninfected cohorts and were analyzed by flow cytometry. The gating strategy for identifying total monocytes, total lymphocytes, myeloid dendritic cells (mDCs), and plasmacytoid dendritic cells (pDCs) is shown in Figure S1. We found that the percentage of total circulating monocytes (relative to total PBMCs) was lower in convalescent COVID-19 patients (23.7%) compared to uninfected controls (38.5%) (Fig. 1a). Conversely, the percentage of lymphocytes was increased in the convalescent population (70% in COVID-19 patients vs. 54.5% in controls) (Fig. 1b). There was no significant difference in the percentages of mDCs or pDCs between the two populations (Fig. 1c-d). The increase in lymphocytes in the convalescent population was driven by an increase in T cells, without a significant difference in the percentages of B cells between the two populations (Fig. 2). There was a slight, but significant, decrease in the percentage of NK cells in convalescent COVID-19 patients (Fig. 2c). These findings are consistent with prior studies demonstrating a decrease in circulating monocytes during convalescent COVID-19 infection ^25, 26^.

**Fig. 1.**
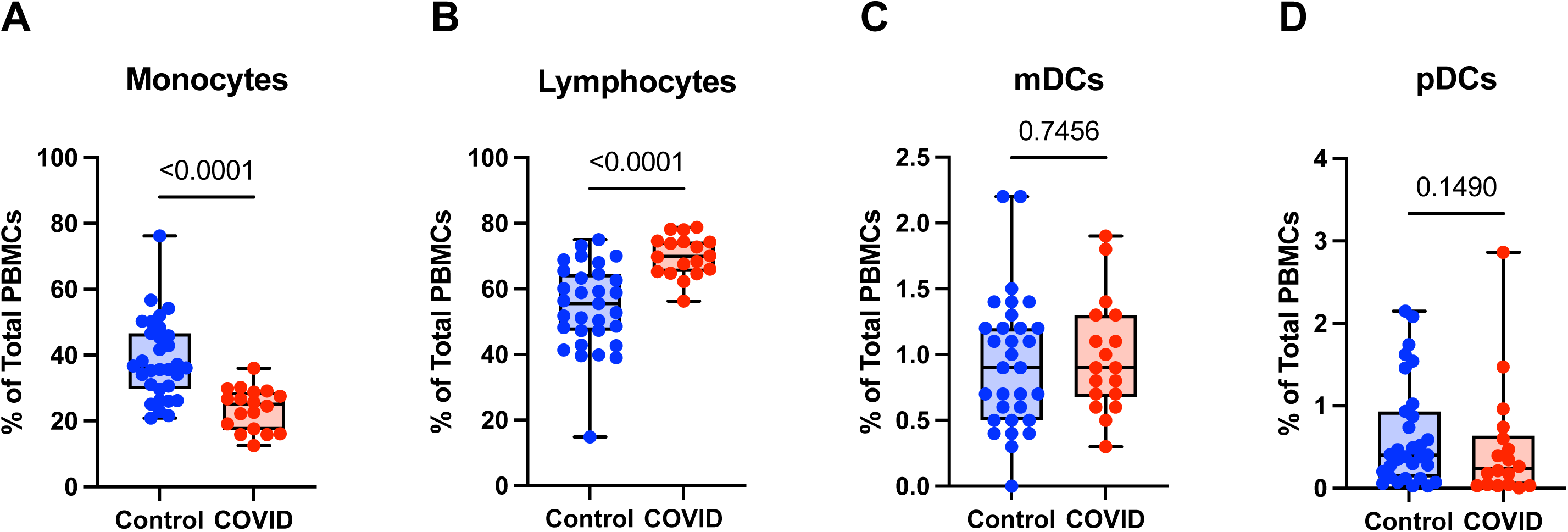
Monocytes are decreased and lymphocytes increased in convalescent COVID-19 patients. (A) Total monocyte percentage, (B) Total lymphocyte percentage, (C) Myeloid dendritic cell percentage, and (D) Plasmacytoid dendritic cell percentage (of PBMCs) in convalescent COVID-19 (blue) and control (red) groups. Mann-Whitney U test. ns, not significant.

**Fig. 2.**
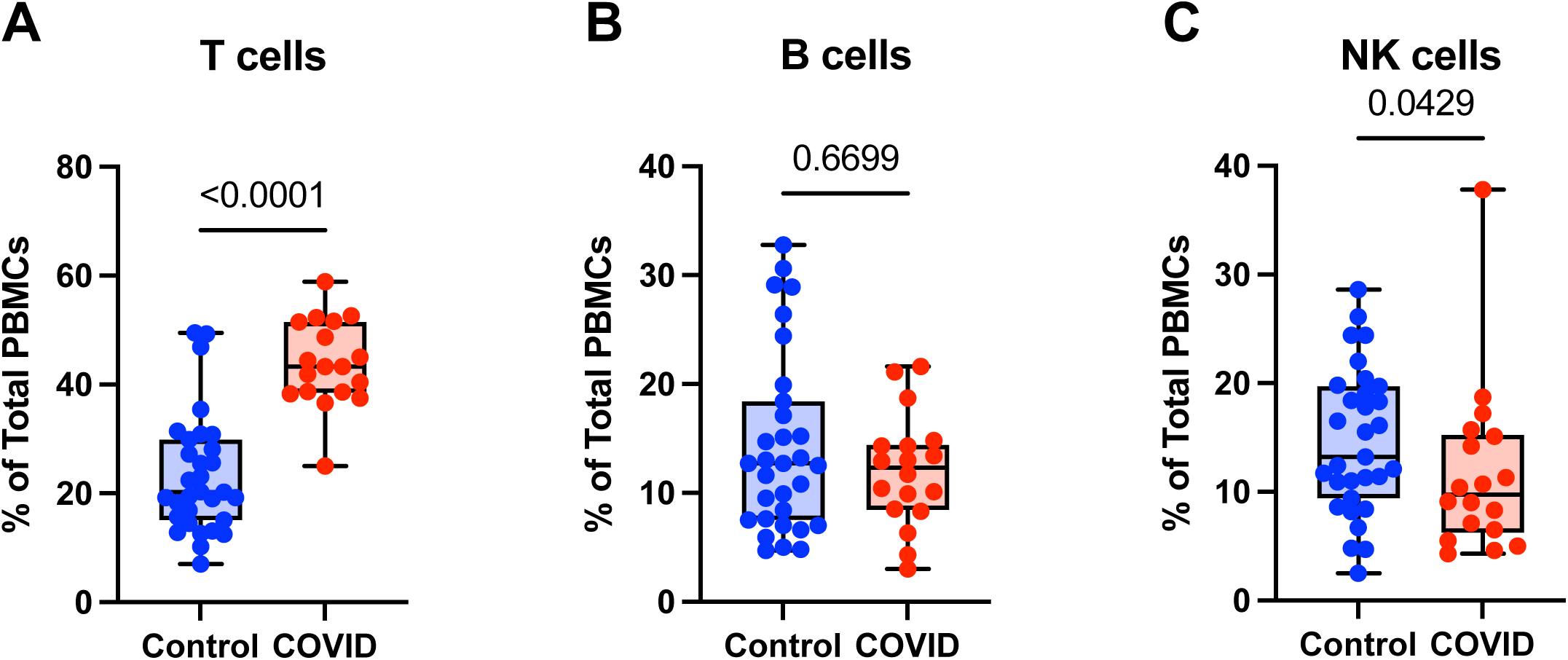
T cells are increased in convalescent COVID-19 patients. (A) T cell percentage, (B) B cell percentage, and (C) NK cell percentage (of PBMCs) in convalescent COVID-19 (blue) and control (red) groups. Mann-Whitney U test. ns, not significant.

Monocyte subsets including classical monocytes (CD14+ CD16-), intermediate monocytes (CD14+ CD16+), and non-classical (CD14^lo^ CD16+) were defined by CD14 and CD16 expression within the total monocyte population (Fig. S1). Whereas the number of classical monocytes was unchanged between the uninfected and convalescent groups (Fig. 3a), the percentage of intermediate and non-classical monocytes were significantly decreased in convalescent patients compared to uninfected donors (2.8% vs. 3.9% and 1.5% vs. 2.6%, respectively) (Fig. 3b-c). Together, these observations suggest that circulating monocyte percentages decrease in convalescent patients, with significant decreases in intermediate and non-classical monocytes.

**Fig. 3.**
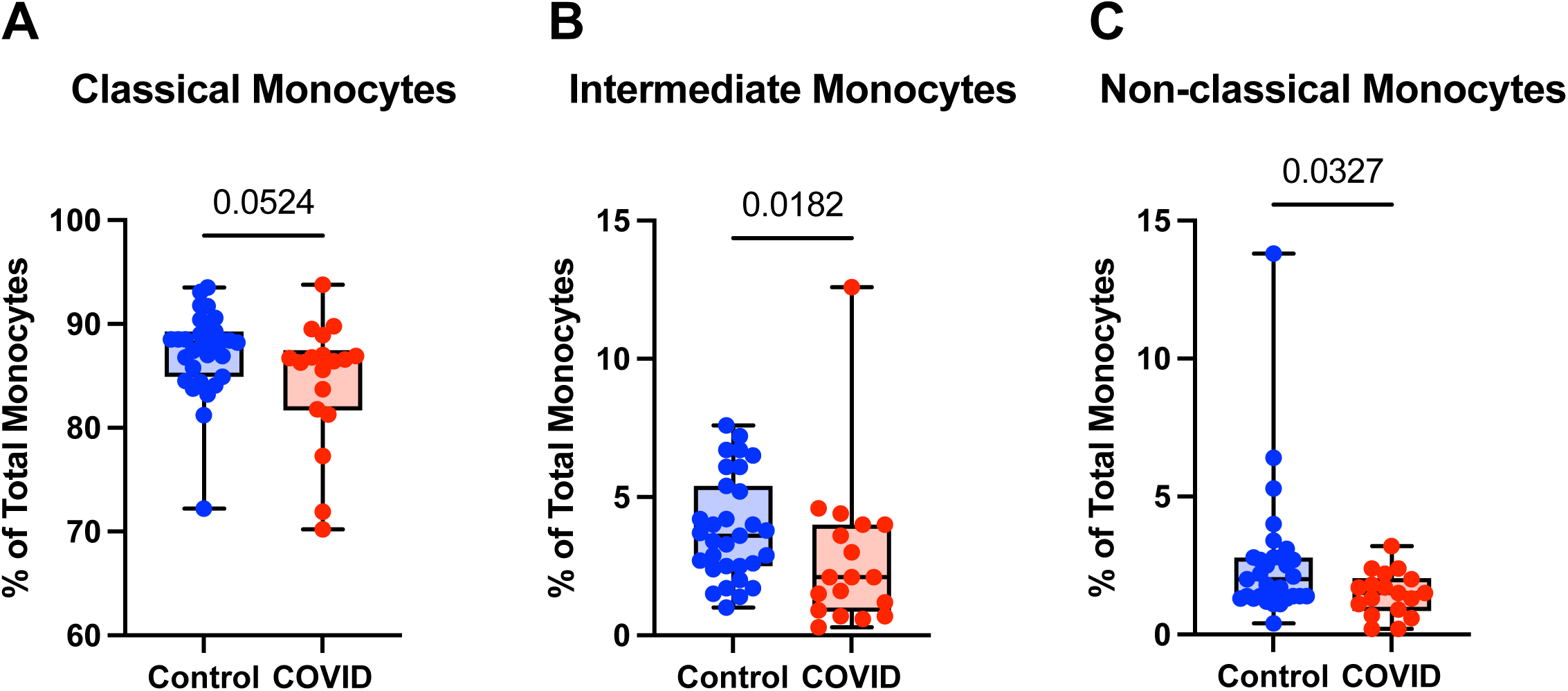
Intermediate and non-classical monocytes are decreased in convalescent COVID-19 patients. (A) Classical monocyte percentage, (B) Intermediate monocyte percentage, and (C) Non-classical monocyte percentage (of total monocytes) in convalescent COVID-19 (blue) and control (red) groups. Mann-Whitney U test. ns, not significant.

### Expression of monocyte activation markers is altered in convalescent COVID-19 patients

To further investigate differences in monocyte activation between the study populations, we assessed the expression of a number of activation markers including: CD4, a glycoprotein expressed at low levels on the surface of monocytes which may stimulate cytokine expression and differentiation to macrophages ^37^ and which is downregulated in response to LPS stimulation ^38, 39^; CD56, an adhesion molecule that is expressed at high levels on monocytes in inflammatory conditions ^40–42^, including severe COVID-19 infection ^43, 44^; CD69, a type II C-lectin receptor upregulated on monocytes in response to stimulation ^45^; CD83, a transmembrane protein involved in the regulation of immune responses ^46^; and CD86, a co-stimulatory molecule expressed in monocytes whose expression can be altered by LPS treatment ^47^. The gating strategy for identifying total monocytes and monocyte subsets for the examination of activation markers is shown in 189 Figure S2.

When comparing classical monocytes from convalescent COVID-19 patients and uninfected controls, we found no significant differences in the percentages of cells expressing bright (T-cell intensity) CD4, CD56, CD69, CD83, or CD86 in the absence of *E. coli* LPS stimulation (Fig. 4). The expression patterns of these markers observed in response to LPS stimulation in these two populations were more complex. In response to stimulation with LPS, we found a statistically significant increase in the percentage of classical monocytes expressing CD56 and CD83, a significant decrease in the percentage of classical monocytes expressing CD69 and CD86, and no change in the percentage of classical monocytes expressing bright CD4 in uninfected control subjects (Fig. 4). In contrast, in convalescent COVID-19 patients, LPS treatment led to a statistically significant increase in the percentage of classical monocytes expressing bright (T-cell intensity) CD4 and CD83, a decrease in the percentage of classical monocytes expressing CD86, and no change in the percentage of classical monocytes expressing CD56 or CD69 from convalescent COVID-19 patients in response to LPS (Fig. 4). When directly comparing LPS-stimulated classical monocytes from convalescent COVID-19 patients to those from uninfected subjects, there was a statistically significant increase in the percentage of cells expressing bright CD4 and CD69, a significant decrease in the percentage of cells expressing CD56, and no difference in the percentage of cells expressing CD83 or CD86. Taken together, these findings suggest that classical monocytes from convalescent COVID-19 patients are hyporesponsive to stimulation with LPS.

**Fig. 4.**
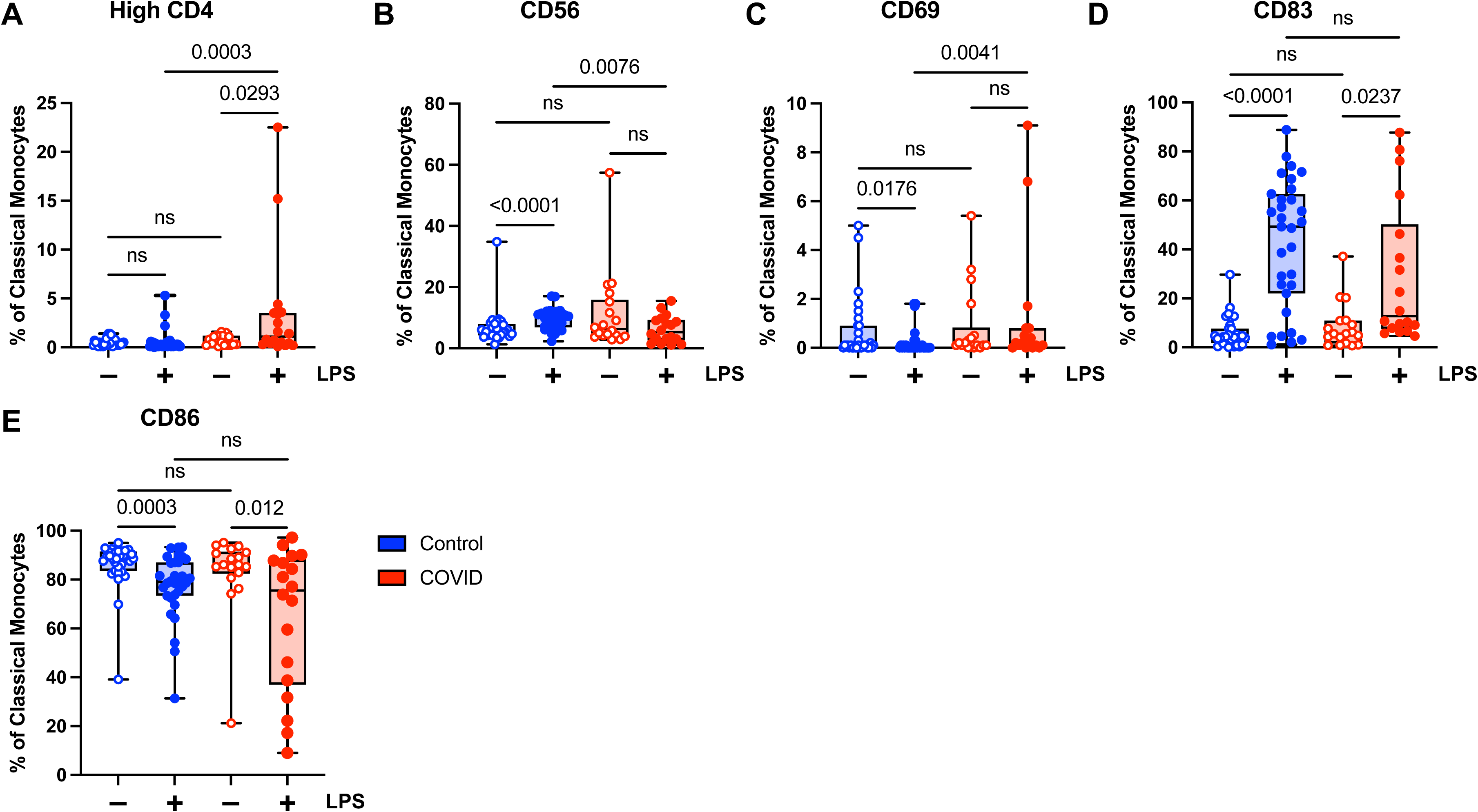
Expression of activation markers is altered in classical (CD14+ CD16-) monocytes from convalescent COVID-19 patients. (A) CD4 percentage, (B) CD56 percentage, (C) CD69 percentage, (D) CD83 percentage, and (E) CD86 percentage (of classical monocytes) in convalescent COVID-19 (blue) and control (red) groups in the absence (empty) or presence (filled) of 100 ng/mL LPS. Comparison between groups (control vs. COVID-19) Mann-Whitney U test. Comparison within groups (untreated vs. LPS-treated) Wilcoxon ranked test. ns, not significant.

Unlike classical monocytes, which showed a complex pattern of activation marker expression, intermediate monocytes demonstrated uniformly decreased expression of bright CD4, CD56, CD69, CD83, and CD86 in response to stimulation with LPS (Fig. 5). This was true for both convalescent and uninfected subjects. In addition, there were no significant differences in any of the activation markers tested when comparing intermediate monocytes from convalescent and uninfected populations, either at baseline or with LPS stimulation (Fig. 5). The differences in activation marker expression between classical and intermediate monocytes in response to LPS stimulation is somewhat surprising given that both monocyte populations express CD14, which is necessary for binding LPS ^48^. These observed differences may result from differential expression of other components of LPS signaling, such as MD2 and TLR4, or downstream effector molecules.

**Fig. 5.**
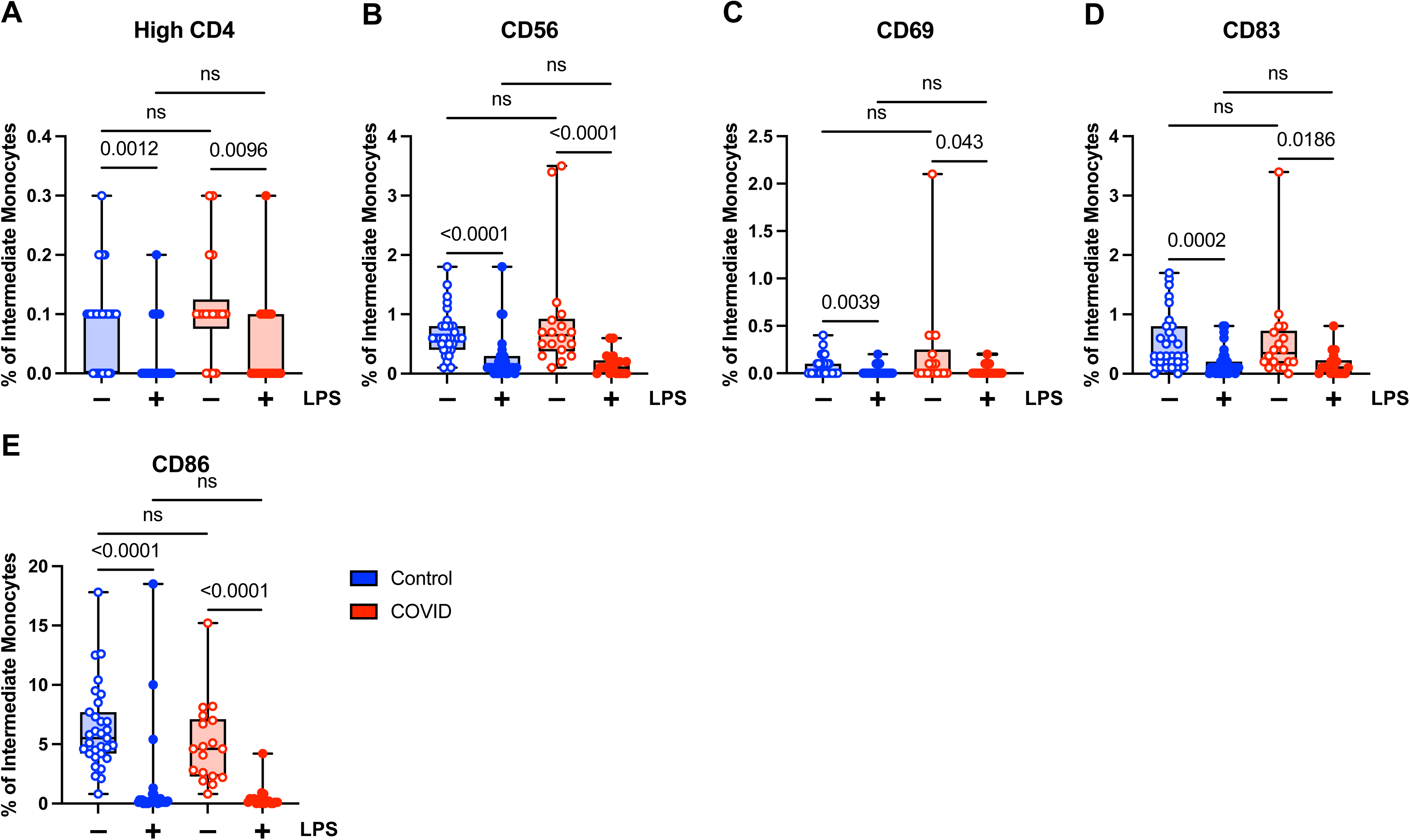
Activation marker expression is not altered in intermediate (CD14+ CD16+) monocytes from convalescent COVID-19 patients. (A) CD4 percentage, (B) CD56 percentage, (C) CD69 percentage, (D) CD83 percentage, and (E) CD86 percentage (of intermediate monocytes) in convalescent COVID-19 (blue) and control (red) groups in the absence (empty) or presence (filled) of 100 ng/mL LPS. Comparison between groups (control vs. COVID-19) Mann-Whitney U test. Comparison within groups (untreated vs. LPS-treated) Wilcoxon ranked test. ns, not significant.

In contrast to classical and intermediate monocytes, which showed no difference in baseline expression of CD56, a higher percentage of non-classical monocytes from convalescent patients expressed CD56 compared to those from uninfected subjects in the absence of LPS stimulation (Fig. S3b). This was the only significant difference in non-classical monocytes between the two study populations. Non-classical monocytes from both convalescent COVID-19 patients and uninfected controls demonstrated decreased CD56 and CD86 expression in response to LPS treatment (Fig. S3b, S3e). In addition, the percentage of non-classical monocytes expressing bright CD4 from uninfected control subjects decreased in response to LPS (Fig. S3a) and the percentage of non-classical monocytes expressing CD83 from convalescent patients decreased in response to LPS treatment (Fig. S3d) The muted response of non-classical monocytes to LPS stimulation is not entirely surprising given that, by definition, these cells express low levels of surface CD14; however, it is possible that the stimulatory effects of LPS could be potentiated by soluble CD14 present in the total monocyte culture.

### Expression of CD142 (tissue factor) is decreased in convalescent COVID-19 patients in response to LPS stimulation

The extrinsic coagulation cascade can be triggered by soluble or cell-surface CD142 (tissue factor). Prior studies have demonstrated increased CD142 expression in the setting of viral infections ^49–52^ and in response to LPS ^49, 53^. In addition, there is a correlation between higher levels of CD142 and COVID-19 disease severity ^51, 52^. Given the role of coagulopathy in COVID-19 pathogenesis, we wanted to evaluate the expression of CD142 by the various monocyte subsets in our uninfected and convalescent COVID-19 populations. We found that although convalescent subjects had a higher percentage of classical monocytes expressing CD142 compared to uninfected subjects (mean 0.68% vs. 0.32%, p=0.0064) at baseline, they had a significantly lower percentage of CD142-expressing classical monocytes in response to LPS treatment (7.2% vs. 10.1%) (Fig. 6a). This was expected given our findings demonstrating decreased activation marker expression in classical monocytes from convalescent COVID-19 patients in response to LPS stimulation (Fig. 4b-c). No differences in CD142 expression were observed between intermediate monocytes from convalescent COVID-19 patients and uninfected controls (Fig. 6b). Similar to our findings with other activation markers, this suggests that classical monocytes from convalescent subjects are less responsive to LPS stimulation.

**Fig. 6.**
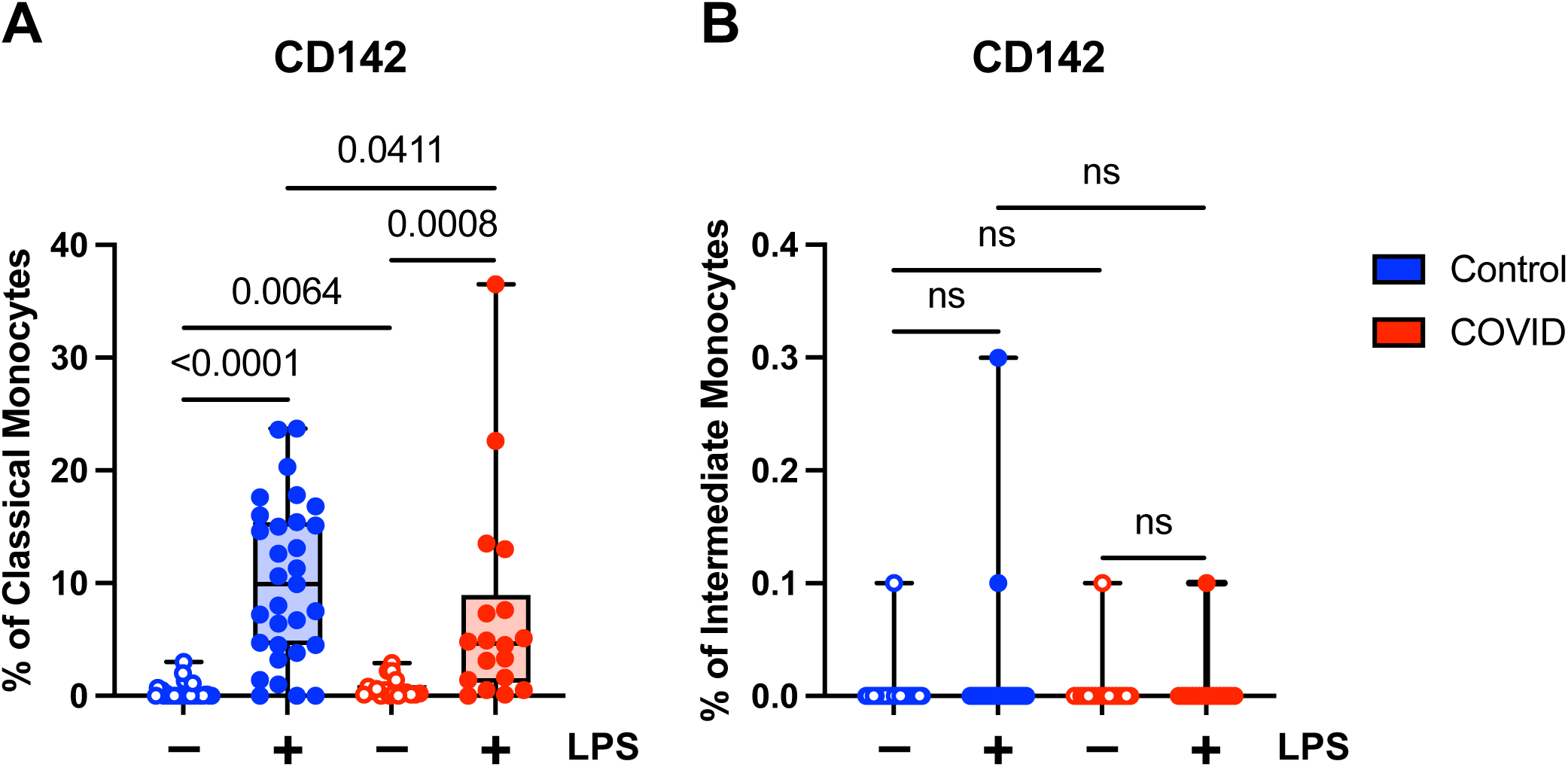
Classical (CD14+ CD16-) monocytes from convalescent COVID-19 patients express less CD142 (tissue factor) in response to LPS. (A) CD142 percentage (of classical monocytes) in convalescent COVID-19 (blue) and control (red) groups in the absence (empty) or presence (filled) of 100 ng/mL LPS. (B) CD142 percentage (of intermediate monocytes) in convalescent COVID-19 (blue) and control (red) groups in the absence (empty) or presence (filled) of 100 ng/mL LPS. Comparison between groups (control vs. COVID-19) Mann-Whitney U test. Comparison within groups (untreated vs. LPS-treated) Wilcoxon ranked test. ns, not significant.

### Expression of proinflammatory cytokines is decreased in convalescent COVID-19 patients

Finally, we wanted to determine whether there were functional differences between monocytes in convalescent COVID-19 patients and uninfected controls. It has been shown previously that monocytes from acutely infected patients produce higher levels of proinflammatory cytokines and type I IFNs ^14–18^. Similarly, monocytes in convalescent individuals have been shown to express increased levels of inflammatory genes ^21–24^. There have been conflicting reports as to whether long COVID is associated with increased levels of proinflammatory cytokines ^32^ ^33–35^ or whether it is characterized by cytokine deficiencies ^36^. Given these disparate study results, as well as our findings suggesting a hyporesponsive monocyte phenotype in convalescent COVID-19 patients, we wanted to determine whether monocytes from convalescent COVID-19 patients were capable of producing proinflammatory cytokines in response to stimuli. We treated purified total monocytes from uninfected and convalescent COVID-19 subjects with LPS and measured TNF-α and IL-6 production by ELISA. We found that monocytes from convalescent COVID-19 patients expressed significantly lower levels of both TNF-α (Fig. 7a) and IL-6 (Fig. 7b) in response to LPS stimulation, consistent with our findings regarding the expression of activation markers (Fig. 4) and cellular CD142 (Fig. 6). In addition, unstimulated monocytes from convalescent patients expressed less IL-6 than those from uninfected controls. Taken together, our findings suggest that monocytes from convalescent COVID-19 patients are hyporesponsive to LPS stimulation.

**Fig. 7.**
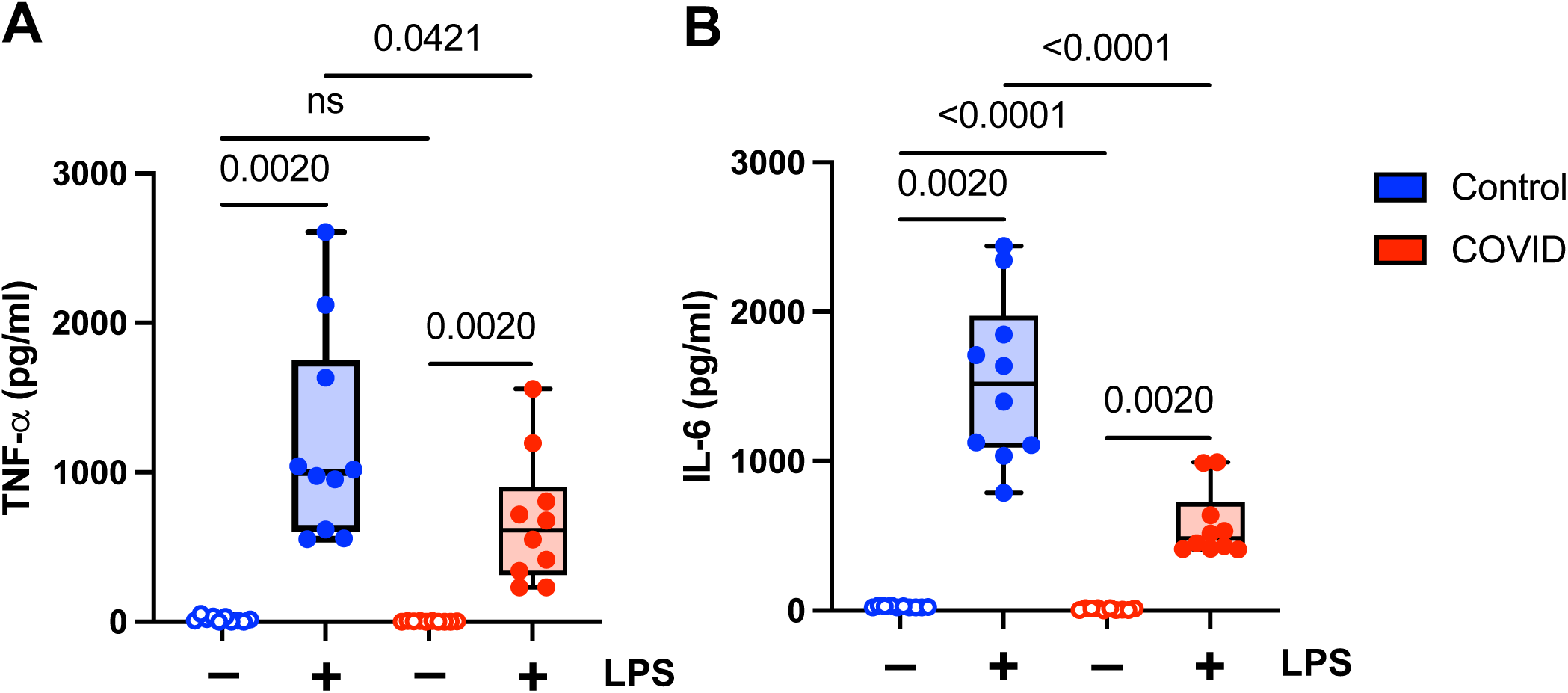
Proinflammatory cytokine expression is decreased in monocytes from convalescent COVID-19 patients. (A) TNF-α and (B) IL-6 expression in convalescent COVID-19 (blue) and control (red) groups in the absence (empty) or presence (filled) of 100 ng/mL LPS. Comparison between groups (control vs. COVID-19) Mann-Whitney U test. Comparison within groups (untreated vs. LPS-treated) Wilcoxon ranked test. ns, not significant.

## DISCUSSION

Several studies examining peripheral blood from COVID-19 patients demonstrated a complex immune response in which myeloid cells play a central role ^54, 55^. The contributions of monocytes to disease pathogenesis varies depending upon the stage of infection, and relatively little is known about their role during the convalescent stage of disease. We aimed to comprehensively analyze the phenotypic and functional changes of monocytes in convalescent COVID-19 patients with an eye toward the potential implications for disease progression.

We show that the percentage of total monocytes is decreased in COVID-19 patients, with significant decreases in intermediate and non-classical monocytes. These results contrast sharply with a previously published study by Park et al. that demonstrated increased circulating total monocytes and increased classical and intermediate monocytes ^21^. The differences between our study and the previously published study by Park et al. may be due to differences in the study populations (primarily Caucasian vs. primarily Asian/Pacific Islander), timing (2-4 weeks post resolution of symptoms vs. >30 days after infection), vaccination status (not vaccinated vs. predominantly vaccinated), or possibly viral strain.

Circulating monocytes are comprised of three phenotypically and functionally distinct subsets: classical monocytes, intermediate monocytes, and non-classical ^56, 57^. Classical monocytes account for about 80–95% of circulating monocytes. These cells are highly phagocytic, and play major roles in innate immune responses, antibody-dependent cellular cytotoxicity (ADCC), and coagulation. Intermediate monocytes account for 2–10% of circulating monocytes. These cells are important producers of reactive oxygen species (ROS), in addition to processing and presenting antigen for T-cell stimulation. Non-classical monocytes account for about 2–10% of circulating monocytes. They are primarily responsible for clearance of debris and apoptotic cells, but also secrete inflammatory cytokines. We found that there were significant decreases in the relative levels of intermediate and non-classical monocytes in convalescent COVID-19 patients. Given their purported functions, their relative decrease may be associated with decreased T-cell activation and/or decreased proinflammatory cytokine production in convalescent patients.

In addition to finding that the percentage of monocytes was decreased in convalescent patients, we demonstrated that they were hyporesponsive to LPS stimulation. Classical monocytes from convalescent COVID-19 patients demonstrate significantly lower levels of CD56 expression in response to LPS stimulation compared to uninfected controls. This finding was not observed in intermediate or classical monocytes. In addition, classical monocytes from convalescent patients show no change in CD69 expression in response to LPS stimulation. Furthermore, monocytes from convalescent patients express significantly less TNF-α and IL-6 when treated with LPS. These findings suggest that, as patients transition from acute to convalescent stages of disease, monocytes shift towards a hyporesponsive phenotype. It is possible that this shift may play a role in dampening the inflammatory response and promoting tissue repair.

Tolerogenic myeloid cells, in particular mDCs, have been shown to suppress inflammation in other settings by means of decreased T-cell activation ^58^. Our findings in convalescent patients are similar to those reported in a cohort of long COVID patients ^36^. In that study, PBMCs from long COVID patients produced lower levels of proinflammatory cytokines compared to uninfected controls.

There is a correlation between expression of cell surface ^51^ and extracellular, vesicular ^52^ CD142 (tissue factor) and COVID-19 disease severity. Interestingly, in a study by Stephenson et al., circulating monocytes displaying the highest levels of ISG expression were found to express ligands and receptors for platelets ^59^. This may explain the increased propensity for coagulopathy in acutely ill COVID-19 patients. We found a significant decrease in classical monocytes expressing CD142 in response to LPS stimulation in convalescent patients compared to uninfected subjects. This finding is consistent with our data showing that classical monocytes from convalescent COVID-19 patients demonstrate significantly lower levels of CD56 and proinflammatory cytokines in response to LPS stimulation.

Our results indicate that there is a relative decrease in monocytes in convalescent COVID-19 patients following mild disease compared to control, uninfected subjects. Intermediate and non-classical monocytes are decreased in COVID-19 patients with no significant difference in classical monocytes. Furthermore, classical monocytes from COVID-19 patients demonstrate decreased CD56 expression and decreased CD142 expression in response to LPS stimulation compared to control subjects. Finally, monocytes from convalescent COVID-19 patients demonstrate significantly decreased TNF-α and IL-6 expression in response to LPS stimulation compared to control subjects. Taken together, these findings suggest that SARS-CoV-2 infection exhibits a clear effect on both monocyte subset development and the response to physiological stimuli, as reflected in their relative numbers, activation state, and function, which lasts into the convalescent phase of disease.

## ACKNOWLEDGEMENTS

This work was supported by funding from ARUP Laboratories Institute for Clinical and Experimental Pathology, the Department of Pathology at the Spencer Fox Eccles School of Medicine, University of Utah (TH), the University of Utah Inflammation, Immunology, and Infectious Disease (3i) Initiative’s Emerging Infectious Diseases Fellowship (ESCPW), a seed grant from the University of Utah Vice President for Research and the Immunology, Inflammation, and Infectious Disease (3i) Initiative (VP), and NIH GRANT 5R01-AI143567 (VP).

## AUTHOR CONTRIBUTIONS

All authors contributed to the study conception and design. Material preparation was performed by Elizabeth Williams, Adam Spivak, and Timothy Hanley. Data collection and analysis were performed by Eugene Ravkov and Timothy Hanley. The first draft of the manuscript was written by Eugene Ravkov and Timothy Hanley. All authors commented on previous versions of the manuscript. All authors read and approved the final manuscript.

## DISCLOSURE OF CONFLICTS OF INTEREST

The authors have no relevant financial or non-financial interests to disclose.

---

**Fig. S1.**
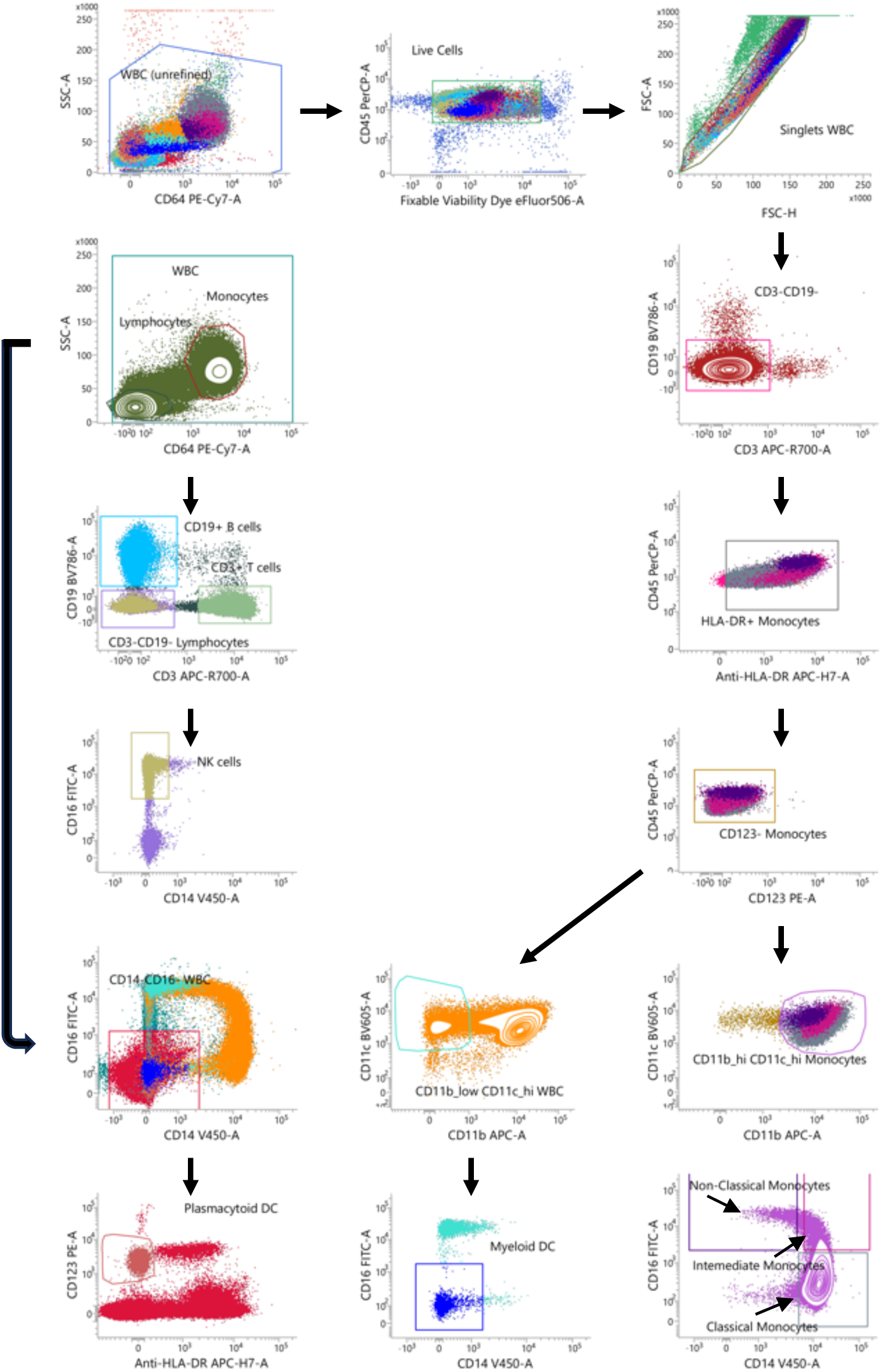
Flow cytometry gating strategy for identification of monocytes, lymphocytes, myeloid dendritic cells (mDCs), and plasmacytoid dendritic cells (pDCs).

**Fig. S2.**
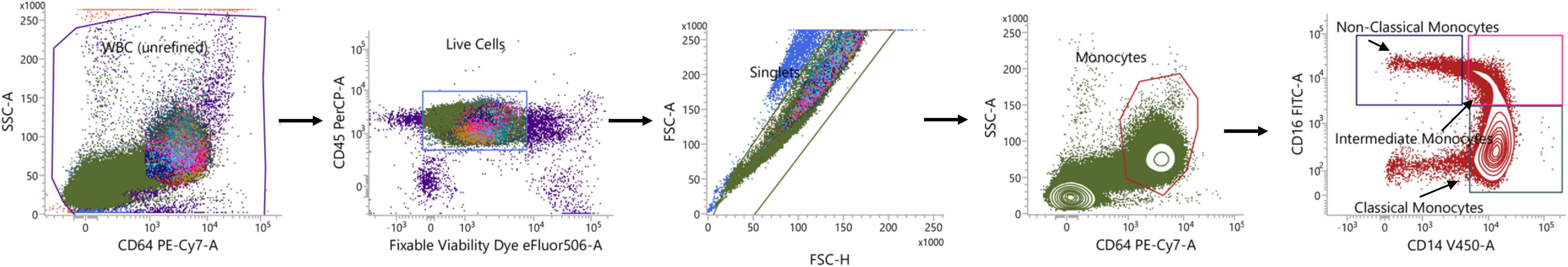
Flow cytometry gating strategy for evaluation of monocyte activation markers in monocyte subsets.

**Fig. S3.**
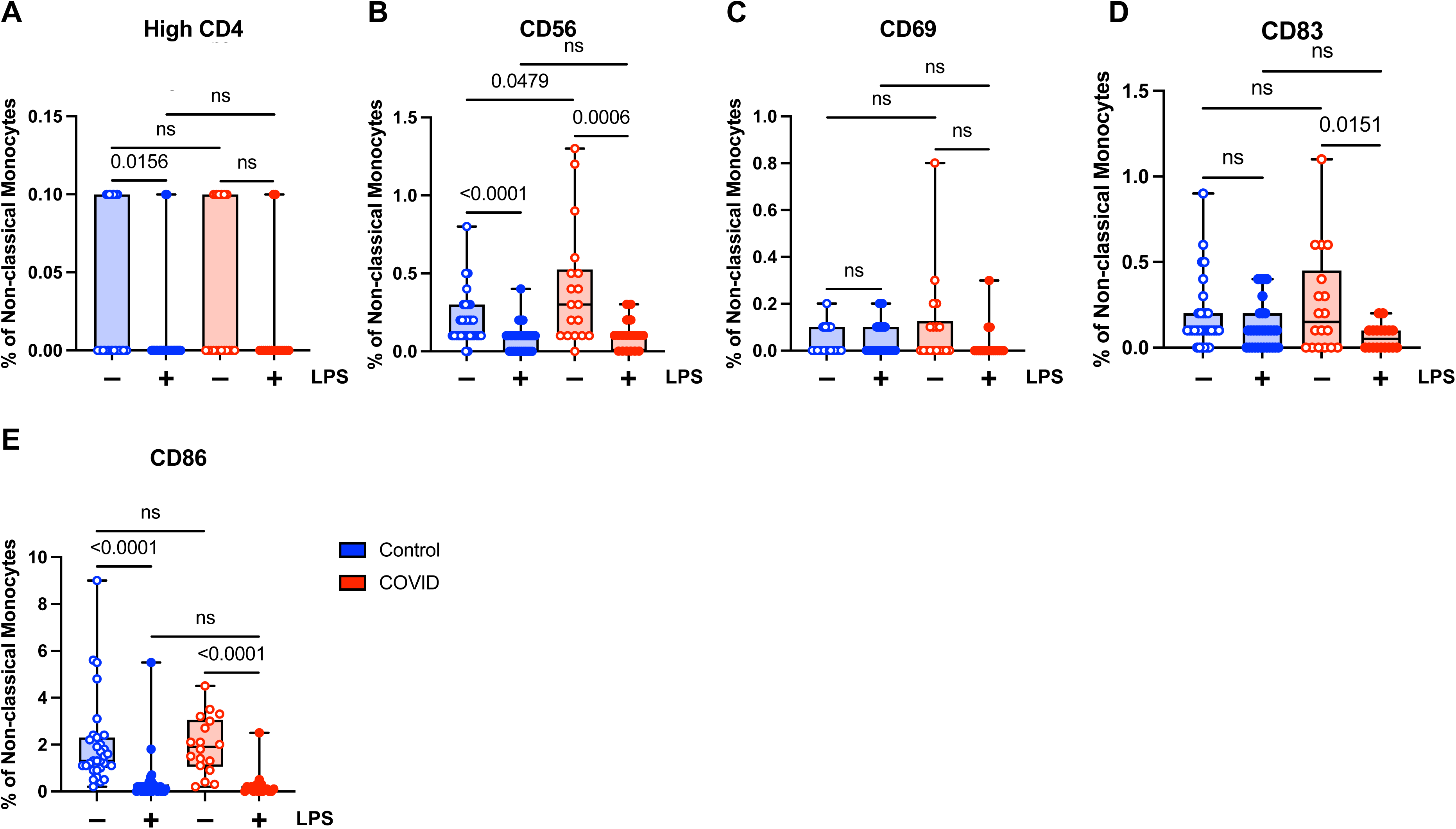
Activation marker expression in non-classical (CD14^lo^ CD16+) monocytes. (A) CD4 percentage, (B) CD56 percentage, (C) CD69 percentage, (D) CD83 percentage, and (E) CD86 percentage (of non-classical monocytes) in convalescent COVID-19 (blue) and control (red) groups in the absence (empty) or presence (filled) of 100 ng/mL LPS. Comparison between groups (control vs. COVID-19) Mann-Whitney U test. Comparison within groups (untreated vs. LPS-treated) Wilcoxon ranked test. ns, not significant.

## Notes

### Competing Interest Statement

The authors have declared no competing interest.

